# Breast Cancer Subtyping with HyperCLSA: A Hypergraph Contrastive Learning Pipeline for Multi-Omics Data Integration

**DOI:** 10.1101/2025.09.22.677517

**Authors:** Gaurav Bhole, HC Poorvi, JR Madhav, P K Vinod, Prabhakar Bhimalapuram

**Author notes:** These authors contributed equally to this work. {, }.

## Abstract

Accurate molecular subtyping of cancer is crucial for advancing personalized medicine. Although multiomics data contain valuable predictive information, effectively integrating them is challenging due to differences across modalities, high dimensionality, and complex cross-modal biological interactions. We propose HyperCLSA (Hyper-graph Contrastive Learning with Self-Attention), a novel deep learning framework for efficient multi-omics integration in breast cancer subtyping. HyperCLSA combines hypergraph-based sample encoding, supervised contrastive learning for latent space alignment, and multi-head self-attention for adaptive fusion of omics modalities. Evaluated on The Cancer Genome Atlas Breast Cancer dataset (TCGA-BRCA) for PAM50 subtype classification, HyperCLSA achieves a state-of-the-art accuracy of 90.1%, significantly outperforming established baselines. Our results demonstrate HyperCLSA’s effectiveness in extracting complementary information across heterogeneous omics sources, providing a robust framework for molecular characterization of breast cancer.

## 1 Introduction

Breast cancer remains the leading cause of cancer-related mortality among women worldwide, with over 2 million new cases diagnosed annually [1]. This disease is characterized by remarkable molecular heterogeneity, which significantly impacts treatment efficiency and patient outcomes [2]. To address this heterogeneity, the PAM50 gene expression assay has been developed to classify breast tumors into intrinsic molecular subtypes—Luminal A, Luminal B, HER2-enriched, Basal-like, and Normal-like—each exhibiting distinct clinical behaviors, treatment responses, and survival patterns [3,4]. Recent advances in high-throughput sequencing technologies have created unprecedented opportunities to examine the disease at multiple molecular levels simultaneously. The integration of diverse omics data types—including genomics, transcriptomics, and proteomics—offers a more comprehensive understanding of breast cancer biology than any single data modality alone.

Existing multi-omics integration methodologies suffer many limitations that compromise their effectiveness. These include high dimensionality, technical noise, inter-modality heterogeneity and the need to capture subtle cross-modal relationships [5]. Addressing these challenges is therefore essential for translating the power of multi-omics data into improving breast cancer subtyping.

Graph-based integration techniques, including specialized Graph Neural Networks like MOGONET [6], have improved performance by modeling inter-sample relationships. However, these methods typically capture only pairwise interactions, failing to represent the higher-order relationships that exist in complex biological systems. While recent frameworks like MORE [7] incorporate hype-graph structures, they have two key limitations. First, their self-attention mechanisms operate on heterogeneous feature spaces without proper normalization, making cross-modal integration less effective. Second, they lack explicit learning objectives that would encourage more discriminative representations across modalities. Overhead in computational time due to uncertainty estimation and interpretability of these models is still a common problem [8]. These shortcomings hinder optimal alignment between different omics data types potentially concealing important biological signals and limiting the discovery of clinically relevant molecular patterns.

To address these challenges in breast cancer subtyping, we introduce **HyperCLSA (Hypergraph Contrastive Learning with Latent Self-Attention)**, a novel three-stage framework for multi-omics integration. First, HyperCLSA constructs modality-specific hypergraph encodings for mRNA, methylation, and miRNA data to capture complex higher-order relationships within each omics view and enrich intra-modal structural representations. Next, supervised contrastive learning aligns these embeddings into a shared latent space, pulling together representations of the same sample across modalities while pushing apart different classes. Finally, a multi-head self-attention mechanism adaptively fuses the aligned embeddings by dynamically weighting each modality’s contribution for every sample. Our evaluation of HyperCLSA on the TCGA-BRCA dataset for PAM50 subtype classification demonstrates its superior performance over strong baselines, showcasing its ability to effectively harness complementary multi-omics information.

## 2 Datasets

This study utilizes two multi-omics datasets: TCGA-BRCA for breast cancer subtyping, comprising 850 samples across five molecular subtypes: Normal-like (44), Basal-like (137), HER2-enriched (52), Luminal A (457), and Luminal B (160) samples. The dataset includes matched mRNA expression, DNA methylation (Illumina HumanMethylation450), and miRNA expression data for each sample.

## 3 Method: The HyperCLSA Pipeline

HyperCLSA is a multi-stage pipeline designed for robust multi-omics integration and classification, as shown in Figure 1.

**Fig. 1.**
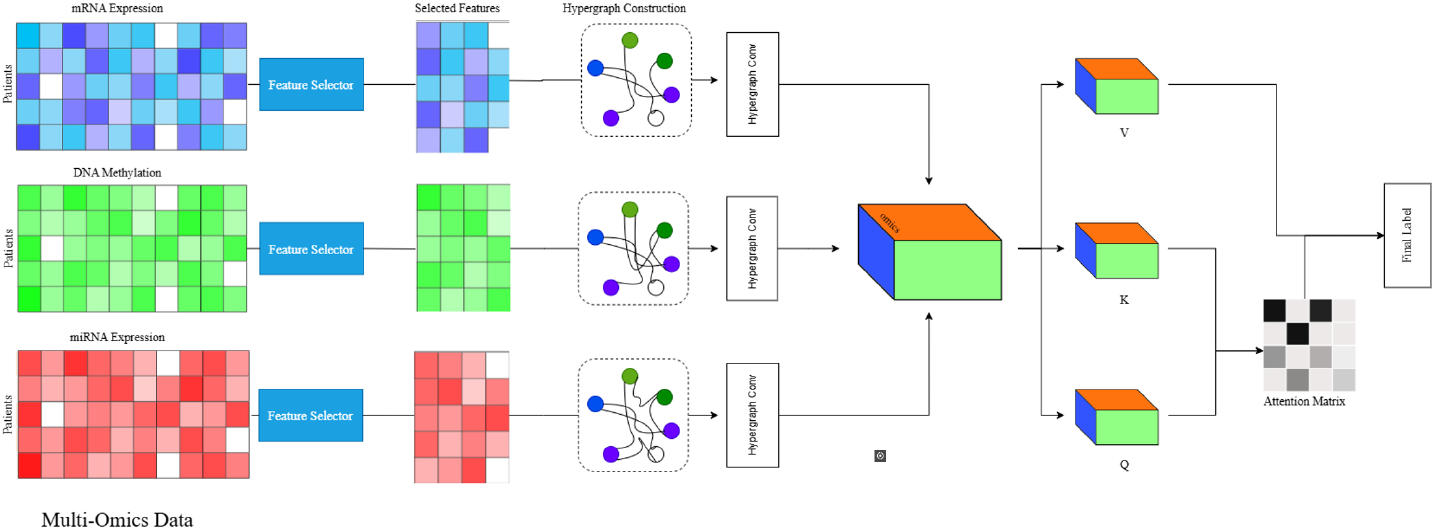
HyperCLSA architecture: modality-specific hypergraph encoding, contrastive alignment, and attention-based fusion

### 1. Feature Selection and Hypergraph Construction

For each of *M* omics views, the raw feature matrix 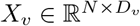 (*N* samples, *D*_*v*_ features) undergoes feature selection. We primarily used Boruta [9], which is an all-relevant feature selection method based on Random Forests. The feature selection is performed independently on each of the views and 5 fold cross-validation is performed to obtain a reduced feature matrix 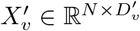.

Subsequently, a hypergraph *H*_*v*_ is constructed for each 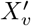 where each sample is a node. We explored k-Nearest Neighbors (k-NN), radius-based, and mutual k-NN methods for defining hyperedges. In the radius-based approach, the hyperedges connect samples whose pair-wise Euclidean distance is below a fixed threshold *ϵ*. For all folds, we fixed *ϵ* at the value that maximized 5-fold CV accuracy in a preliminary grid search. The resulting hypergraph is represented by an incidence matrix 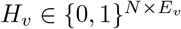 (where *E*_*v*_ is the number of hyperedges, often *N* if centered on each node), from which a normalized hypergraph Laplacian is computed, facilitating information propagation in the HGNN encoder [10,11].

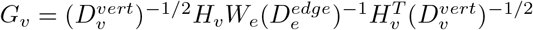

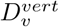 and 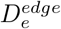 are diagonal matrices of vertex and hyperedge degrees, respectively, and *W*_*e*_ contains hyperedge weights (typically identity).

### 2. View-Specific Encoding and Latent Space Alignment

Each omics view *X*_*v*_ is processed through a dedicated two-layer Hypergraph Convolutional Network (HGNN). The network computes embeddings as 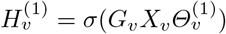 and 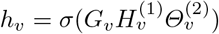, where *G*_*v*_ is the hypergraph Laplacian, *Θ* values are learnable parameters, and *σ* is the LeakyReLU activation. These view-specific embeddings are then projected into a common latent space using *z*_*v*_ = Linear_*v*_(*h*_*v*_) with L2-normalization. To encourage samples of the same class to cluster together across different omics views, we apply a supervised contrastive loss:

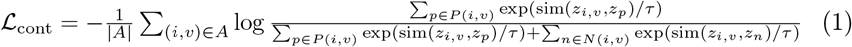

where *P* (*i, v*) contains embeddings from the same class as sample *i, N* (*i, v*) contains embeddings from different classes, sim(*·, ·*) is cosine similarity, and *τ* is a temperature parameter. This approach aligns the latent representations while preserving class distinctions.

### 3. Multi-Head Attention for View Integration

For each sample *j*, we integrate information across the normalized view embeddings *{z*_*j*,1_, …, *z*_*j,M*_*}* using multi-head self-attention. Self attention is computed over the embeddings and multiple attention heads capture different relationship patterns between views. Outputs are then concatenated and linearly projected to produce 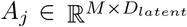. The final fused representation *z*_*fused,j*_ is calculated by averaging across views. This representation is passed through a classification layer to obtain subtype probabilities. The model optimizes a combined objective:

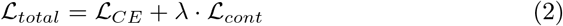

where ℒ_*CE*_ is the cross-entropy loss and *λ* balances the contribution of the contrastive term.

## 4 Experiments and Results

### 4.1 Experimental Setup

We evaluated HyperCLSA on the PAM50 subtype classification task using The Cancer Genome Atlas Breast Cancer (TCGA-BRCA) dataset. This dataset provided matched mRNA expression (RNA-Seq FPKM-UQ), DNA methylation (Illumina HumanMethylation450 BeadChip beta values), and miRNA expression (miRNA-Seq RPM) data for 850 samples across five molecular subtypes. PAM50 subtype labels were obtained from the TCGA Pan-Cancer Clinical Data Resource (TCGA-CDR) annotations.

### Data Preprocessing and Feature Selection

For each omics modality, features with low variance (standard deviation < 0.1) or high missingness (> 20% of samples) were removed. The remaining features were then standardized using Z-score normalization. Feature selection was performed independently for each modality within each training fold of the cross-validation to prevent data leakage. For the Boruta method, which yielded the best results (Table 2, input data for each view was first imputed using mean imputation and then scaled using `StandardScaler`. Boruta was then applied using a Random Forest estimator (100 trees, `max_depth=5`, balanced class weights) and configured to run for a maximum of 50 iterations.

### Hypergraph Construction

Following feature selection, modality-specific hypergraphs were constructed. In the optimal radius-based approach, hyperedges were formed by connecting samples if their pairwise Euclidean distance was below a defined threshold *ϵ*. This threshold *ϵ* was dynamically determined for each omics view within each training fold as the median of all pairwise sample Euclidean distances calculated on the selected features of the training samples. A normalized hypergraph Laplacian *G*_*v*_ was then computed for each view to facilitate information propagation in the HGNN encoder.

### Model Configuration and Training

Each omics view’s selected features were input into a dedicated two-layer Hypergraph Convolutional Network (HGNN) encoder with hidden layer dimensions of [400, 256]. These encoders utilized `LeakyReLU` activation functions and a dropout rate of 0.5 after each activation. The resulting view-specific embeddings were then projected into a common latent space of 64 dimensions using a linear layer followed by L2-normalization. The multi-head self-attention mechanism responsible for fusing these latent representations employed a single attention head with a dropout rate of 0.3 on the attention scores and output. The HyperCLSA model was trained using the Adam optimizer, where training proceeded for a maximum of 5000 epochs, with an early stopping criterion (patience of 200 epochs) based on the validation loss. The total loss (Eq. 2) function combined a standard cross-entropy term for classification (*L*_*CE*_) with a supervised contrastive loss term (*L*_*cont*_).

### Evaluation Protocol and Baselines

Model performance was evaluated using accuracy, macro-averaged F1-score (F1-macro), and weighted-averaged F1-score (F1-weighted). A 5-fold stratified cross-validation protocol was employed, and results are reported as mean *±* standard deviation across the folds. For comparison, we benchmarked HyperCLSA against several baselines: (1) MOGONET, a graph convolutional network approach (2) MORE, a hypergraph-based neural network method; and (3) HyperTMO, a hypergraph-based convolutional network without any attention strategies for multi-omics data.

### 4.2 Main Performance Results

Table 1 provides a summary of classification performance, comparing HyperCLSA with state-of-the-art baselines on the TCGA-BRCA dataset. HyperCLSA, configured with Boruta feature selection and radius-based hypergraph construction demonstrated superior performance, significantly outperforming MOGONET, MORE and HyperTMO.

**Table 1.** Performance Comparison on TCGA-BRCA for PAM50 Subtype Classification (5-fold CV mean ± std). HyperCLSA results are for its best configuration (Boruta FS, Radius Graph, all 3 omics)

| Method | Accuracy | F1-macro | F1-weighted |
| --- | --- | --- | --- |
| MOGONET | 0.829±0.018 | 0.774±0.017 | 0.825±0.016 |
| MORE | 0.835±0.020 | 0.768±0.021 | 0.820±0.023 |
| HyperTMO | 0.858±0.023 | 0.821±0.019 | 0.863±0.023 |
| **HyperCLSA** | **0.901±0.007** | **0.866±0.019** | **0.901±0.007** |

**Table 2.** Ablation Study: Impact of Feature Selection on HyperCLSA (BRCA, Radius Graph, All 3 Omics)

| Feature Selection (with Radius Graph) | Accuracy | F1-macro | F1-weighted |
| --- | --- | --- | --- |
| None | 0.795±0.022 | 0.672±0.040 | 0.784 ±0.023 |
| RFE | 0.852±0.019 | 0.795±0.024 | 0.847±0.020 |
| Boruta | **0.901±0.007** | **0.866±0.019** | **0.901±0.007** |

### 4.3 Ablation Studies

To assess the contribution of key components, we performed the ablations: **Impact of Feature Selection (FS):** Table 2 shows results using radius-based hypergraph generation. Boruta FS significantly outperformed Recursive Feature Elimination (RFE) and a baseline without any FS. This underscores the critical role of effective feature selection in reducing noise and dimensionality.

#### Impact of Hypergraph Construction Method

Table 3 shows results using Boruta FS. Radius-based hypergraph construction yielded the best performance, surpassing mutual k-NN and standard k-NN. This highlights the sensitivity of model performance to the method chosen for capturing inter-sample relationships. An additional ablation study on different attention mechanisms given in Table S1 confirmed that HyperCLSA’s multi-head self-attention provided benefits over simpler aggregation methods.

**Table 3.** Ablation Study: Impact of Hypergraph Construction Method on HyperCLSA (BRCA, with Boruta FS, All 3 Omics)

| Hypergraph Method (with Boruta FS) | Accuracy | F1-macro | F1-weighted |
| --- | --- | --- | --- |
| k-NN | 0.851±0.020 | 0.795±0.036 | 0.848±0.021 |
| Mutual k-NN | 0.871±0.020 | 0.840±0.020 | 0.872±0.018 |
| Radius-based | **0.901±0.007** | **0.866±0.019** | **0.901±0.007** |

#### Impact of Using Different Omics

Using all three omics modalities together yields the best classification performance, as shown in Fig 2. Among the individual modalities, mRNA emerges as the most informative single view. Incorporating either miRNA or methylation alongside mRNA largely maintains this high level of performance. In contrast, excluding mRNA—or relying solely on a non-mRNA modality—leads to the most significant drop in accuracy. These results confirm the complementary value of integrating all three omics types.

**Fig. 2.**
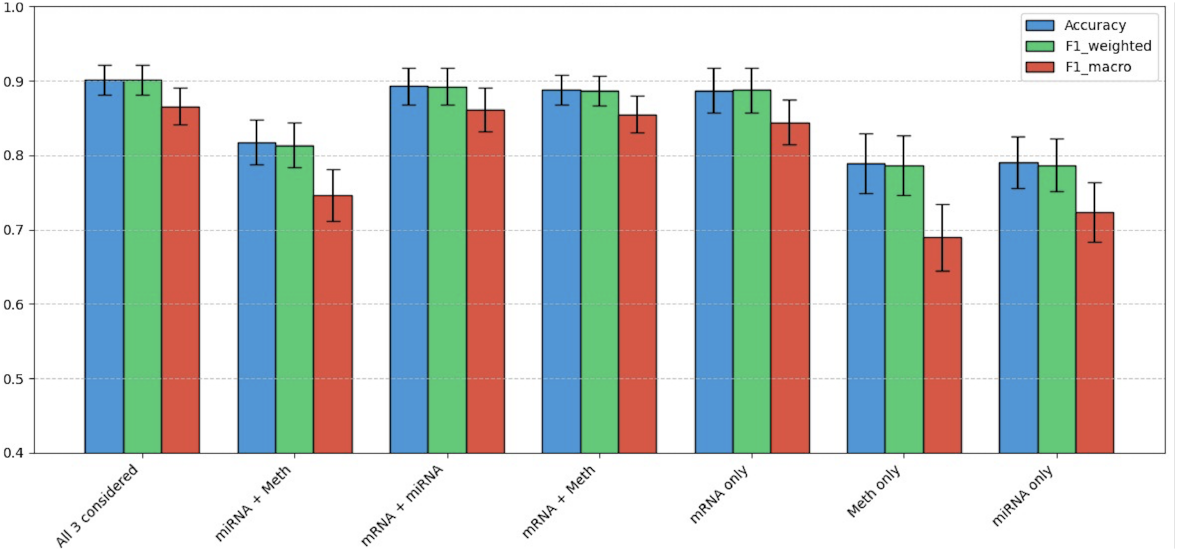
Ablation: Performance of HyperCLSA using different omics combinations on TCGA-BRCA. All three modalities yield the best results

## 5 Discussion

HyperCLSA demonstrates robust performance in breast cancer PAM50 subtyping by effectively integrating mRNA, DNA methylation, and miRNA data. The PAM50 intrinsic subtypes were originally defined primarily based on gene expression (mRNA) profiles [12]. This explains the strong performance observed when using mRNA data alone with HyperCLSA (Fig. 2). Despite being a graph-based method, our implementation is highly efficient and scalable, requiring only 2GB of GPU memory on the TCGA-BRCA dataset, suggesting that HyperCLSA is scalable to larger datasets using standard computational infrastructure.

However, breast cancer is a complex disease driven by alterations across multiple molecular layers. We can observe from Fig. 2 that integrating additional omics modalities consistently enhances classification performance. The combination of all three omics types (mRNA, DNA methylation, and miRNA) yielded the highest accuracy (0.901) and F1-macro (0.866). This improvement, while modest over mRNA-only given PAM50’s origin, signifies that DNA methylation and miRNA data provide complementary, biologically relevant information. DNA methylation plays a crucial role in gene regulation, and aberrant methylation patterns are well-documented drivers in breast cancer, often contributing to subtype-specific gene silencing or activation [13,14]. miRNAs, as post-transcriptional regulators, also show dysregulation patterns correlated with breast cancer subtypes and progression [15]. HyperCLSA’s self-attention mechanism adaptively weighs diverse signals for more robust classification, especially on ambiguous samples. This synergy is evident as multi-omics combinations, such as miRNA with any of the other omics, significantly outperform their single-omic counterparts.

Although demonstrated on breast cancer, the HyperCLSA pipeline is designed to be broadly applicable across cancer types. Its extension to other cancers could be promising but is currently limited by the availability of large, high-quality multi-omics datasets with well-annotated subtypes. Key limitations of the current work include challenges in model explainability and the need for careful hyperparameter tuning. Future work will focus on enhancing biological interpretability. For instance, by applying post-hoc explanation techniques like SHAP (SHapley Additive exPlanations) to the trained model, we can move beyond aggregate performance metrics. This would allow for the identification of key driving features and omics modalities for each individual prediction, potentially revealing patient-specific biomarkers and offering deeper biological insights into the model’s classification decisions. Additionally, incorporating more biologically informed hypergraph constructions, for example using curated cancer-specific pathway data to refine hyperedges, represents a promising avenue, though the development and integration of such resources remains ongoing.

## 6 Conclusion

This paper introduced HyperCLSA, a novel deep learning pipeline that effectively integrates multi-omics data for precise breast cancer subtype classification. By combining view-specific hypergraph encoding, utilizing optimal strategies like Boruta for feature selection, leveraging radius-based hypergraph construction, supervised contrastive learning for robust latent space alignment, and multi-head self-attention for adaptive fusion of complementary information, HyperCLSA achieves state-of-the-art PAM50 subtyping performance on the TCGA-BRCA dataset. Our results highlight the importance of integrating mRNA, DNA methylation, and miRNA data and demonstrate the robustness of the proposed architectural components. HyperCLSA marks a significant advancement toward more accurate patient stratification in breast cancer and offers a flexible framework with potential applications across precision oncology.

## Data Availability and Code

All TCGA-BRCA data used are publicly available from the Broad GDAC Firehose portal. The code for HyperCLSA is available at: https://github.com/Gaurav2543/HyperCLSA.

