## Supplementary material for "Breast Cancer Subtyping with HyperCLSA: A Hypergraph Contrastive Learning Pipeline for Multi-Omics Data Integration": Table S1

### S1: Detailed Implementation Details

The HyperCLSA model was implemented using PyTorch (Version 1.13) and Python (Version 3.9). For the TCGA-BRCA dataset, we utilized three omics views: mRNA expression (RNA-Seq FPKM-UQ), DNA methylation (Illumina HumanMethylation450 BeadChip beta values), and miRNA expression (miRNA-Seq RPM). Data was preprocessed by filtering features with low variance (std $< 0.1$) or high missingness ($>$20% samples), followed by Z-score standardization per feature. PAM50 subtype labels (Luminal A, Luminal B, Basal-like, HER2-enriched, Normal-like) were obtained using TCGA Pan-Cancer Clinical Data Resource (TCGA-CDR) annotations, which are based on standard PAM50 classifiers.

Training was performed using the Adam optimizer with a learning rate of $5 \times 10^{-4}$ and weight decay of $1 \times 10^{-5}$. The model was trained for a maximum of 5000 epochs, with early stopping if the validation loss did not improve for 200 consecutive epochs. A batch size of 128 was used where applicable (though for graph-based methods on the full dataset, batching applies differently).

The HGNN encoders consisted of two layers with hidden dimensions [400, 256]. The shared latent space dimension $D_{latent}$ was set to 64. The multi-head self-attention mechanism employed $h = 1$ attention head, with a dropout of 0.3 applied to the attention scores and the output of the attention layer. The dropout rate within the HGNN encoders (after activation functions) was 0.5. The hyperparameter $\lambda_{contrast}$ balancing the classification and contrastive losses was set to 0.295. The temperature $\tau$ for the contrastive loss (Eq. 1) was 0.1.

For hypergraph construction:

- **k-NN**: $k = 6$ nearest neighbors were used based on Euclidean distance. Each sample and its $k$ neighbors formed a hyperedge.
- **Radius-based**: Hyperedges connected samples within a Euclidean distance $\epsilon$. The radius $\epsilon$ was chosen as the median of all pairwise sample distances within each omics view after feature selection, calculated per fold.
- **Mutual k-NN**: Similar to k-NN with $k = 6$, but an edge (and thus hyperedge participation) between two samples $i$ and $j$ required $j$ to be in $i$'s k-NN list and $i$ to be in $j$'s k-NN list.

The incidence matrix $H_v$ was constructed such that $H_{ji} = 1$ if sample $j$ is part of the hyperedge centered at sample $i$. Hyperedge weights in $W_e$ were set to identity.

Feature selection parameters:

- **RFE**: Used with a logistic regression estimator. Aimed to select approximately 1000 features for mRNA and methylation, and around 500 for miRNA, per fold. The exact number varied slightly based on CV performance during RFE's internal steps.

– **Boruta**: Used with a Random Forest estimator (100 trees, max_depth=5). The number of selected features was determined by the algorithm based on Z-scores of feature importance compared to shadow features.

All experiments were conducted using a 5-fold stratified cross-validation scheme, ensuring class proportions were maintained in train/test splits. Feature selection and hypergraph construction were performed strictly within the training data of each fold to prevent data leakage.

### S2: Additional Ablation Study Details

Table S1 shows the impact of different attention/fusion mechanisms explored during development, along with different feature selectors. These "Model Variants" refer to earlier exploratory architectures. "Baseline Fusion" often implies a simpler concatenation or weighted sum without explicit cross-modal attention. "Cross-Modal Attention" usually refers to transformer-like attention between different modality embeddings before fusion. "No-Attention (Simple Aggregation)" typically means averaging or concatenating latent embeddings without any attention mechanism. "Pathway Guided Attention" was an experiment to weigh attention based on pathway information, which proved complex to tune effectively without curated pathway lists. The results for "No-Attention (Simple Aggregation)" compared to the final HyperCLSA self-attention highlight the benefit of the sophisticated fusion used.

**Table S1.** Supplementary Ablation: Impact of Attention Variants and Feature Selection on HyperCLSA (BRCA, All 3 Omics). Model variants refer to earlier explorations not part of the final HyperCLSA self-attention architecture described in the main paper, but serve to illustrate the importance of sophisticated fusion.

| Model Variant (Exploratory) | Feat. Sel. | Accuracy | F1-macro | F1-weighted |
|---|---|---|---|---|
| Baseline Fusion (No X-attn) | RFE | 0.84 | 0.83 | 0.84 |
| X-Modal Attention | RFE | 0.81 | 0.81 | 0.82 |
| No-Attention (Aggr.) | RFE | 0.79 | 0.79 | 0.81 |
| Pathway Guided Attn | RFE | 0.83 | 0.82 | 0.84 |
| Baseline Fusion (No X-attn) | Boruta | 0.82 | 0.81 | 0.82 |
| X-Modal Attention | Boruta | 0.83 | 0.82 | 0.84 |
| No-Attention (Aggr.) | Boruta | 0.77 | 0.76 | 0.77 |
| Pathway Guided Attn | Boruta | 0.82 | 0.81 | 0.82 |

The results show that combining all three omics (mRNA, miRNA, and methylation) achieves the best performance, with accuracy of 90.1% ± 2.0%, F1-weighted of 90.1% ± 2.0%, and F1-macro of 86.6% ± 2.5%. Among individual modalities, mRNA alone performs the best (88.7% accuracy), which is expected since PAM50 subtype classification is based on mRNA expression. Removing mRNA leads to a notable drop in performance, while combining mRNA with either miRNA or methylation retains high accuracy (88–89%). Single-omic models using only miRNA or methylation perform the worst (79%), highlighting the complementary value of integrating multiple omics.

### S3: Further Discussion on Pathway Integration and Generalizability

**Pathway-Informed Hypergraphs:** Our initial attempt to enhance prediction accuracy by incorporating pathway knowledge into the hypergraph construction module utilized the Reactome Database. This database provides mappings between Gene Ensemble IDs and pathways. For each pathway, we generated a hyperedge where the distance between samples was governed exclusively by genes belonging to that pathway, subject to thresholding (e.g., only samples with significant alteration in that pathway were connected). However, the inclusion of all pathways from the Reactome database (numbering in the thousands) introduced significant noise. This is because the majority of cataloged pathways may not exhibit substantial changes during breast cancer progression or might not be relevant for distinguishing PAM50 subtypes. This highlights the necessity of subsetting only cancer-specific or, more precisely, breast-cancer-subtype-relevant pathways for analysis. The current lack of comprehensive, easily accessible, and machine-readable online resources that catalog cancer-specific pathways specifically curated for different omics data types presents a challenge. We hypothesize that the performance of our model could be substantially improved if this pathway integration were refined to focus on a curated set of biologically relevant pathways, underscoring the need for specialized cancer pathway databases that are also structured for computational integration.

**Generalizability to Other Cancers:** While this study focused on breast cancer due to the well-established PAM50 subtypes and relatively good data availability in TCGA, the HyperCLSA pipeline is designed with a modular and generic architecture. Its core components—view-specific feature selection, hypergraph-based encoding for higher-order relationships, contrastive learning for inter-modal alignment, and adaptive attention-based fusion—are, in principle, applicable to multi-omics datasets from other cancer types or even other complex diseases.

The primary challenge for broader application is often data-related:

1. **Data Availability and Quality:** Many other cancer types may not have datasets as comprehensive as TCGA-BRCA in terms of sample size, number of matched omics modalities per sample, and quality of data. Smaller sample sizes can make it harder to train complex deep learning models and can also make hypergraph construction less stable.
2. **Subtype Definitions:** Subtype definitions for other cancers might be less standardized or biologically distinct compared to the PAM50 subtypes in breast cancer. The effectiveness of supervised contrastive learning and classification depends heavily on the quality and relevance of the ground truth labels.

3. **Biological Heterogeneity:** Different cancer types are driven by different molecular mechanisms. While the pipeline is generic, the optimal feature sets, the nature of inter-sample relationships (and thus optimal hypergraph construction parameters), and the relative importance of different omics modalities might vary significantly. Therefore, adapting HyperCLSA to a new cancer type would likely require:
   - Re-optimization of feature selection strategies (e.g., the number of features to select, or even the choice between RFE and Boruta might change).
   - Tuning of hypergraph construction parameters (e.g., $k$ in k-NN, or $\epsilon$ in radius-based methods).
   - Potentially adjusting model hyperparameters like latent dimensions or attention head numbers.

Despite these challenges, the fundamental principles of learning robust, aligned representations from multiple data sources and adaptively fusing them are valuable across diverse oncological contexts. HyperCLSA provides a flexible framework that can be adapted and tuned for such scenarios, provided sufficient data and relevant biological context are available.
